# A precision medicine approach for *HCN1* Developmental and Epileptic Encephalopathy

**DOI:** 10.1101/2024.01.09.574555

**Authors:** Lauren E. Bleakley, Chaseley E. McKenzie, Da Zhao, Ming S. Soh, James Spyrou, Ian C. Forster, Bang V. Bui, Christopher A. Reid

## Abstract

Pathogenic variants in *HCN1* causing cation leak result in a severe developmental and epileptic encephalopathy (DEE). Current treatment options for patients with *HCN1*-DEE are limited and are insufficient to fully address both the seizures and clinical comorbidities of this disorder.

Org 34167 is a brain penetrant broad-spectrum HCN channel inhibitor that has completed phase I clinical trials. We used a range of assays at molecular, cellular, network and behavioural levels to explore the potential of Org 34167 as a precision medicine for *HCN1*-DEE.

Org 34167 restored the voltage sensitivity of the DEE HCN1^M305L^ mutated channel, significantly reducing cation leak. It also restored I_h_-mediated ‘sag’, hyperpolarised the resting membrane potential and reduced firing of layer V neurons from the Hcn1^M294L^ mouse model of *HCN1*-DEE, which was engineered based on the HCN1^M305L^ pathogenic variant. Additionally, Org 34167 reduced neuronal epileptiform activity and restored retinal light sensitivity in these mice, suggesting it may improve both seizures and other clinical comorbidities. However, Org 34167-mediated tremors were noted at therapeutic doses. Org 34167 was also effective at reducing cation leak caused by five additional *HCN1* pathogenic variants, suggesting broader utility.

Overall, these data demonstrate that a small molecule HCN inhibitor can restore channel and consequent physiological functions, positioning it as a promising precision therapeutic approach for *HCN1*-DEE.

## Introduction

*HCN1* encodes HCN1 channels that play a unique biological role in controlling neuron excitability, including setting the resting membrane potential, controlling synaptic integration, and generating resonance properties.^1^ Pathogenic variation in *HCN1* is an established cause of epilepsy, encompassing syndromes that can range from mild through to very severe.^2–5^ A subset of *HCN1* pathogenic variants cause Developmental and Epileptic Encephalopathy (DEE), a syndrome that is characterised by drug-resistant epilepsy and significant developmental delay.^6^ Although a full genotype-phenotype relationship is yet to be established, *HCN1* pathogenic variants that cause cation leak associate strongly with DEE.^7^ Electrophysiological recordings from pyramidal neurons of the Hcn1^M294L^ heterozygous knock-in mouse (homolog of the HCN1^M305L^ variant) confirm cation leak, with the resulting depolarised resting membrane potential taking neurons closer to their firing threshold, thus increasing excitability.^8^ A similar depolarised neuronal resting potential has been reported in the Hcn1^G380D^ heterozygous mouse model (homolog of HCN1^G391D^).^9^ We predict that the majority of *HCN1*-DEE cases involve variants that result in cation leak.^10^ Therapeutic strategies that target HCN1 channel cation leak are therefore expected to improve outcomes for a high proportion of *HCN1*-DEEs.

Here we explore a small molecule precision approach for *HCN1*-DEE. Org 34167 is a propyl-1,2 benzisoxazole derivative that was initially synthesised by Organon Laboratories for the treatment of depression (Patent WO/1997/040027).^11,12^ Org 34167 is a broad-spectrum modulator of HCN channels.^13^ The impact of Org 34167 on channel function includes a hyperpolarising shift in the voltage dependence of activation and a slowing of activation kinetics.^13,14^ Furthermore, a reduction in the maximum open probability at extreme hyperpolarisation argued for an additional voltage-independent mechanism.^14^ Importantly, Org 34167 is highly brain penetrant.^13^ Org 34167 completed phase I clinical trials and was well tolerated at low doses (<2.5 mg), with only moderate side effects seen at doses as high as 40 mg,^12^ positioning it as a possible repurposing candidate for *HCN1*-DEE.

Here we test the utility of Org 34167 in a preclinical model of *HCN1*-DEE. At a biophysical level, Org 34167 re-established the voltage sensitivity of the HCN1^M305L^ channel, reducing cation leak. Org 34167 also reduced interictal spiking on electrocorticography (ECoG), normalised some behavioural deficits, and recovered retinal function in the Hcn1^M294L^ mouse model of DEE. Biophysical analysis of five additional *HCN1*-DEE variants demonstrated a similar reduction in cation leak suggesting that Org 34167 may be used more broadly in this disease. These preclinical data provide strong motivation to consider Org 34167 as a potential precision therapeutic treating both seizures and other comorbid states in *HCN1*-DEE.

## Materials and methods

### Animal housing and husbandry

Adult female *Xenopus laevis* frogs were housed at the Florey Institute of Neuroscience and Mental Health prior to oocyte extraction. Mice were bred on-site at the Florey Institute of Neuroscience and Mental Health. Male heterozygous Hcn1^M294L^ mice were mated with wild-type C57BL/6J female mice, yielding either Hcn1^+/+^ (Hcn1^WT^) or Hcn1^M294L/+^ (Hcn1^M294L^) offspring. Mice were housed in standard 15 x 30 x 12 cm cages maintained under 12 hour dark and light cycles (<50 lux), with access to dry pellet food and tap water *ad libitum*. For all experiments both male and female mice were used, with littermates used wherever possible. Mice were acclimatised to experimental rooms for at least 30 minutes before experimentation. At the conclusion of experimentation mice were culled by cervical dislocation or decapitation following deep isoflurane anaesthesia, which are ANZCCART approved methods. For both mouse and frog studies, anaesthesia and analgesia were used where appropriate. All animal research reported in this manuscript adheres to Animal Research: Reporting of In Vivo Experiments (ARRIVE) guidelines.

### *Xenopus* oocyte electrophysiology

#### Solutions and reagents

Standard OR-2 solution was used to wash defolliculated oocytes, and contained (in mM) 82.5 NaCl, 2 KCl, 1 MgCl_2_.6H_2_O, 5 HEPES, adjusted to pH 7.4 with TRIS. Injected *Xenopus laevis* oocytes were incubated in a standard ND96 storage solution that contained (in mM): 96 NaCl, 2 KCl, 1 MgCl_2_.6H_2_O, 1.8 CaCl_2_.2H_2_O, 5 HEPES, adjusted to pH 7.4 with TRIS and supplemented with antibiotic gentamicin (50 mg/L). Electrophysiology was performed using a superfusing solution that contained (in mM) 100 KCl, 1.8 BaCl_2_, 1 MgCl_2_, 10 HEPES, pH 7.4 adjusted with TRIS (100K solution). Ba^2+^ was substituted for Ca^2+^ to minimise contamination from endogenous Cl^-^ and K^+^ channel currents.

CsCl was added as powder to the 100K working solution to give a final concentration of 10 mM. Org 34167 was prepared from powder (supplied by SYNthesis medchem, Australia) as a stock concentration in 10% DMSO and water, which was further diluted in the 100K working solution to the test concentration (100 μM). Stock Org 34167 in solution was stored at 4°C and was found to be stable for at least 4 weeks.

#### cRNA preparation

cDNA encoding wild-type human HCN1 (RefSeq NM_021072.4, Ensemble database) and DEE variants (S272P; M305L; I380F; I380N; A387S and G391D) was subcloned into the pGEMHE-MCS vector containing the 5’ and 3’ UTRs from *Xenopus* β-globin to improve its expression in oocytes.^15^ The plasmid sequence was verified by bi-directional Sanger sequencing, then linearised with NheI-HF (New England Biolabs) and purified using QIAquick PCR Purification Kit (QIAGEN). The concentration and quality of the linearised plasmid were confirmed using NanoDrop Spectrophotometer (Thermo Fisher Scientific) and gel electrophoresis. Linearised plasmid was transcribed to cRNA using the T7 mMessage mMachine kit (Ambion) and purified using RNeasy Mini Kit (QIAGEN). NanoDrop Spectrophotometer and gel electrophoresis were used to check the concentration and quality of cRNA, which was stored at −80°C.

#### Expression in Xenopus laevis oocytes

Adult female *Xenopus laevis* frogs were anesthetised with 1.3 mg/mL tricaine methanesulfonate (MS-222) and ovaries surgically removed via a small incision to the abdomen. Oocytes were defolliculated with 1.5 mg/mL collagenase for 2 h. Separated oocytes were washed with calcium-free OR-2 solution and healthy mature oocytes stage V or VI were isolated for experiments. Oocytes were injected manually with 10 ng of cRNA in a 50 nL injection solution and were maintained in ND96 storage solution at 17°C for 2-3 days before experimentation.

#### Two-electrode voltage clamp and voltage step protocols and data analysis

Standard two-electrode voltage clamp hardware was used (TEC-05X or TEC10X, NPI), with series resistance (R_s_) compensation applied to ensure clamp accuracy. Electrodes were filled with 3 M KCl with resistances between 0.2-0.5 MΩ. Voltage clamp control and data acquisition were under software control (pClamp version 10, Molecular Devices). All experiments were performed at 18-20°C. Oocytes were placed in a small perfusion chamber (filled volume ∼20 μL) that was continuously perfused by gravity feed via a common manifold and controlled by pinch valves. Current-voltage *(IV)* relationships were generated using a voltage step protocol with 10 mV steps of 2.5 s duration from a −30 mV holding potential to test potentials in the range −110 mV to +20 mV. Data were sampled at 200 µs/point and low pass filtered at 500 Hz using the voltage clamp inbuilt analog Bessel filter. Cs^+^ is a blocker of HCN channels and was used to distinguish between leak from endogenous channels and HCN channels.^16,17^ All analyses of the co-expressed channels (50% HCN mutant + 50% HCN WT) and contemporaneous WT controls were performed on Cs^+^-subtracted currents.

#### Data analysis

Data were analysed using Prism 3.02 (GraphPad) and Clampfit 10.4 (Molecular Devices). For quantification of steady-state HCN activation, the current induced by voltage steps was taken as the steady-state current measured over an approximately 100 ms interval at end of the test pulse. Cs^+^ subtraction was applied by subtracting post-acquisition data sets with 10 mM CsCl and 100 μM Org 34167 as required. Activation time constants were obtained from exponential fits to unsubtracted data, commencing the fit after the initial inflexion in activation current, using Clampfit routines. For tail current analysis, baseline correction was applied after the current had reached a steady-state and the instantaneous current was measured at the start of the repolarising step (*I*_tail_) immediately after the capacitive transient had settled and fit with a form of the Boltzmann equation:

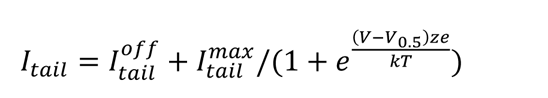

where *I^max^_tail_*is the maximum instantaneous tail current and *I^off^_tail_* is the offset reported by the fit (typically = 0 with baseline correction), *V*_0.5_ is the mid-point voltage, *z* the apparent valence of the charge moved, and *kT/e*=25.3 mV at 20°C. Data sets were then offset corrected and normalised to the predicted *I^max^_tail_* and pooled.

### Whole-cell patch-clamp electrophysiology

Slice electrophysiology studies were conducted on tissue from male and female Hcn1^M294L^ mice (*n* = 5 mice, aged P29-75).

#### Solutions

Brain slices were cut in a chilled cutting solution containing (in mM): 125 choline chloride, 20 D-Glucose, 0.4 CaCl_2_·2H_2_O, 6 MgCl_2_·6H_2_O, 2.5 KCl, 1.25 NaH_2_PO_4_ and 26 NaHCO_3_. Slices were maintained in an artificial cerebrospinal fluid (aCSF) containing (in mM): 125 NaCl, 10 D-Glucose, 2 CaCl_2_·2H_2_O, 2 MgCl_2_·6H_2_O, 2.5 KCl, 1.25 NaH_2_PO_4_ and 26 NaHCO_3_. For all recordings, the electrode internal solution contained (in mM): 125 K-gluconate, 5 KCl, 2 MgCl_2_·6H_2_O, 10 HEPES, 4 ATP-Mg, 0.3 GTP-Na, 10 phosphocreatine, 0.1 EGTA and 0.2% biocytin; with a pH of 7.2.

#### Slicing protocol

Mice were deeply anaesthetised using isoflurane and decapitated. Brains were rapidly removed and mounted in a slice chamber containing chilled cutting solution and bubbled continuously with carbogen gas. 300 µm thick coronal slices were cut using a vibratome (Leica VT1200S). Slices were transferred to a holding chamber filled with aCSF and bubbled continuously with carbogen gas for 30 minutes at 32°C and then at least another 30 minutes at room temperature prior to recording.

#### Recording parameters

Slices were mounted in a recording chamber and continually perfused with aCSF bubbled with carbogen gas (2 mL/min) at a bath temperature of 32°C. Whole-cell patch clamp electrophysiological recordings were made from layer V somatosensory cortical pyramidal neurons (layer V neurons) using borosilicate glass electrodes with initial resistance of 3–7 MΩ. Recordings were made using an Axon Multiclamp 700B amplifier (Molecular Devices), Digidata 1550 digitizer (Molecular Devices), and pCLAMP version 10 software. Data were sampled at 50 kHz with a low-pass filter at 10 kHz. All slice electrophysiology data were analysed using AxoGraph X (version 1.7.6). Recordings were conducted at baseline and following wash-on of 30 µM Org 34167 as outlined below.

#### Slice electrophysiology recordings

All slice electrophysiology data were collected in current clamp mode. Bridge balance and pipette capacitance neutralisation were manually applied. Baseline current injection-action potential (AP) output relationships were established using a step protocol of 2 s square pulses from –100 pA to + 450 pA in 25 pA steps, conducted from rest. Input-output (*i-o*) curves were constructed from raw numbers of events automatically detected in AxoGraph using an amplitude threshold of at least 70 mV. Sag was elicited with a –250 pA current injection from rest. Sag was defined as the difference between the most hyperpolarised point within the first second and the final ‘steady state’ of the voltage trace. To measure the effect of Org 34167 on resting membrane potential, a 5-minute gap-free recording in the absence of holding current was made, during which Org 34167 was applied. Resting membrane potential at baseline was calculated as the average resting membrane potential in the first minute of this trace (prior to Org 34167 application), while resting membrane potential following Org 34167 was calculated as the average resting membrane potential in the final minute of this trace, once Org 34167 has been in the bath for approximately 2 minutes. *I-o* and sag protocols were repeated in the presence of 30 µM Org 34167 in order to measure the effect of the drug on these parameters.

### Electrocorticography (ECoG)

#### ECoG electrode implantation surgery

Male and female Hcn1^M294L^ mice (*n* = 25) were used for ECoG experiments to study the effect of Org 34167 on neuronal epileptiform activity. ECoG electrode implantation was performed at >P24. Mice were anaesthetised via isoflurane inhalation (4% for induction, 1-2% for maintenance throughout surgery), placed on a stereotaxic frame, and administered meloxicam (2.5 mg, subcutaneous, Ilium) as an analgesic and lidocaine hydrochloride (10 mg, subcutaneous, Pfizer) to provide local anaesthesia to the scalp. An incision was made and three holes, each 1 mm in diameter, were drilled into the skull. Two of these holes were drilled bilaterally over the somatosensory cortex, with the third located immediately caudal to the lambdoid suture 0.5 mm lateral from the midline towards the right side of the skull. Stainless steel screw electrodes were implanted into each hole, with the anterior two screws implanted epidurally and used as the active channel electrodes, and the posterior screw implanted slightly more shallowly and used as the reference channel electrode. A ground electrode made of silver wire was affixed to the skull immediately caudal to the lambdoid suture 0.5 mm lateral from the midline towards the left side of the skull. The reference, ground, and two active channel electrodes were connected to a mouse electroencephalography (EEG) head mount (8201-EEG Pinnacle technology Inc.) via silver leads (Cat No. 785500, A-M Systems Inc.) and soldered in place. Self-curing acrylic resin was used to hold the head mount and electrodes in place. Mice were left to recover for at least one week prior to experimentation.

#### ECoG recordings

Mice were placed into individual clear plexiglass containers (140 cm x 175 cm x 140 cm) for the duration of each ECoG recording. The head mount was linked to a mouse EEG preamplifier (8406-SE, Pinnacle Technology Inc.) and connected to a 4-channel data conditioning/acquisition system (8200-K1-SE3, Pinnacle Technology Inc.). ECoG data were sampled at 250 Hz and filtered (40 Hz low-pass, 0.5 Hz high-pass) using Sirenia Acquisition software (version 2.1.0, Pinnacle Technology Inc.). To test the effect of Org 34167 on neuronal activity, baseline ECoG activity was recorded for 2 hours, before mice were administered Org 34167 (1 mg/kg or 2 mg/kg, dissolved in 0.9% w/v saline) or vehicle control intraperitoneally (30G needle), and then ECoG activity was recorded for a further 2 hours. All ECoG recordings were conducted during the light phase of the light-dark cycle.

#### ECoG data analysis

Spike analysis was performed by visual inspection of raw ECoG data visualised and extracted using Sirenia Seizure Pro software (version 1.7.5, Pinnacle Technology Inc.). ‘Spikes’ had distinctive morphology and were defined as biphasic events lasting < 200 ms with at least thrice the amplitude of baseline ECoG activity.^8,10^ Baseline spikes were counted from a 1 hour recording epoch prior to drug administration. Spikes post-Org 34167 were counted from an epoch 10 to 70 minutes following drug administration. Mice were excluded from all analyses if they experienced a spontaneous seizure during baseline recordings. Mice were also excluded if there was significant noise (electrical noise or movement artefacts) present in the recording trace. All spike analyses were completed by a blinded experimenter.

### Behaviour

The effect of Org 34167 on behaviour of Hcn1^M294L^ mice was assessed using the open-field locomotor assay, the elevated plus maze, and the rotarod assay. Male and female mice aged P60-99 took part in behavioural experiments.

#### Open-field locomotor assay

Mice (*n* = 33 Hcn1^WT^, 10 Hcn1^M294L^) were placed in a square 27.3 x 27.3 x 20.3 cm open-field arena and could move freely for one hour, during which time infrared rays tracked their movement. Data indicating distance travelled and other parameters was recorded and compiled using MED Associates Activity Monitor software (Med Associates Inc.). Within each genotype, mice were randomly allocated to receive either 2 mg/kg Org 34167 or vehicle control (saline) via intraperitoneal (i.p.) injection 10 minutes before the assay commenced.

#### Elevated plus maze

The elevated plus maze consists of a plus-shaped platform raised 40 cm from the floor. Two of the arms, on opposite sides, are enclosed with 55 cm high walls extending from the centre, with the other two arms remaining open with no walls. Mice (*n* = 5 Hcn1^WT^, 10 Hcn1^M294L^) were individually placed in the centre of the maze facing one of the open arms and were allowed to freely roam the maze for 10 minutes, and their movements were tracked by a ceiling-mounted video camera. Entries into each arm, along with their respective durations, were measured using tracking software (CleverSys TopScan Lite). Percentage of time spent in the open arm was calculated as

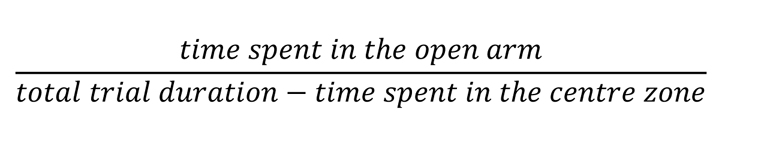

Hcn1^M294L^ mice were randomly allocated to receive either 2 mg/kg Org 34167 or vehicle control (saline) via i.p. injection 10 minutes before their trial. All Hcn1^WT^ mice received saline.

#### Rotarod

Mice (*n* = 10 Hcn1^WT^, 9 Hcn1^M294L^) were trained to walk on the rotarod over three sessions: two 2-minute sessions at a fixed speed of 4 rpm, and one 2-minute session accelerating from 4-20 rpm. Mice were then tested on the rotarod for one 10-minute trial, consisting of 5 minutes where the rotarod accelerated from 4-40 rpm, followed by up to 5 minutes at a fixed speed of 40 rpm. Hcn1^M294L^ mice were randomly allocated to receive either 2 mg/kg Org 34167 or vehicle control (saline) via i.p. injection 10 minutes before this test. All Hcn1^WT^ mice received saline. Latency to fall was recorded for each mouse and analysed.

### Electroretinography (ERG)

A total of 21 Hcn1^M294L^ mice were used. Before the procedure, animals were dark-adapted overnight, then weighed and anaesthetised with an i.p. injection of ketamine (80 mg/kg) and xylazine (10 mg/kg) (Troy Laboratory). The mixture was diluted in sterile injectable saline (1:10) to aid with hydration and ease of administration (10 μL/g). Topical anaesthesia and pupil mydriasis were achieved with single drops of proxymetacaine 0.5% and tropicamide 0.5% (Alcaine™ and Mydriacyl™, respectively, Alcon Laboratories). Corneal hydration was maintained with lubricating eye drops during ERG (Celluvisc, Allergan). During ERG assessment, body temperature was maintained at 37°C to prevent hypothermia. At the end of *in vivo* assessment, anesthetised animals were culled by cervical dislocation.

Experiments were conducted in a lightproof room, and all preparation was undertaken with only the aid of a dim red headlight to minimise light exposure. Animals were set up in a Faraday cage to minimise electromagnetic noise interference. Following induction of anaesthesia and mydriasis, animals were lightly secured to a water heated platform to minimise movement and breathing artifacts. Custom made chlorided silver active electrodes (A&E Metal Merchants) were placed upon the apex of the corneas, with reference ring-shaped electrodes placed around the equator of both eyes. A grounding needle electrode (Grass) was then inserted subcutaneously into the tail, before positioning the Ganzfeld bowl at eye level. Raw signals were processed via a pre-amplifier (P511 AC Amplifier, Grass) and a main amplifier (ML785 Powerlab 8SP, ADInstruments Pty Ltd) with band-pass filter setting of 0.1 to 1000 Hz. Signals were recorded using Scope™ software (ADInstruments) and exported for offline analysis.

A 30G needle was placed into the intraperitoneal space containing either 2 mg/kg Org 34167 or vehicle control (saline). Animals were light-adapted (140 cd/m^2^) for 5 minutes prior to baseline signal acquisition, which comprised a short (2.07 log cd.s/m^2^, each 10 repeats 2 seconds apart) and a long flash (256 ms, 2.37 log cd.s/m^2^). The drug or vehicle was then injected after which the effect was monitored every 2 minutes using the same flashes. After 30 minutes a full photopic intensity response function was acquired (0.76 – 2.67 log cd.s/m^2^) along with responses to progressively higher temporal frequencies at a fixed light intensity (3.99 log cd.s/m^2^ from 5 to 45 Hz in 5 Hz steps, 50% duty cycle).

### Statistical analyses

Statistical analyses were performed using GraphPad Prism software (version 8.1.0–9.2.0). Data were first analysed using a Shapiro-Wilk test for normality. Statistical significance was then determined using a two-tailed Student’s *t*-test (paired or unpaired as appropriate) for normally distributed data, a Wilcoxon test for paired data where the Shapiro-Wilk test returned a result of < 0.05, or a Mann-Whitney U-test for unpaired data where the Shapiro-Wilk test returned a result of < 0.05. ERG data across groups and parameters (i.e. time, stimulus intensity or stimulus frequency) was analysed using 2-way repeated measures ANOVA. All data points are shown as mean ± standard error of the mean (SEM). Statistical significance for all comparisons was set at *p* < 0.05.

### Study approval

All experiments were approved by the Animal Ethics Committee at the Florey Institute of Neuroscience and Mental Health prior to commencement. Animals were monitored in line with protocols approved by this committee. Experiments were performed in accordance with the Prevention of Cruelty to Animals Act, 1986 under the guidelines of the National Health and Medical Research Council (NHMRC) of Australia Code of Practice for the Care and Use of Animals for Experimental Purposes.

### Data availability

Raw data and images are available upon request from the corresponding author.

## Results

### Biophysical analysis of Org 34167 on HCN1^M305L^ channel function

Org 34167 is a brain penetrant propyl-1,2 benzisoxazole derivative that is a broad-spectrum modulator of HCN channels^12,13^ (Fig. 1A, inset). Two-electrode voltage-clamp recordings were obtained from *Xenopus laevis* oocytes to investigate the impact of Org 34167 on the mutant HCN1^M305L^ channel. As previously reported, HCN1^M305L^ causes a dissociation between the voltage sensor and pore domain resulting in a channel that is constitutively open with minimal voltage dependence^8,18^ (Fig. 1A). Remarkably, Org 34167 (100 µM) reinstated some voltage dependence to HCN1^M305L^ channels which suggested that it can partially restore the functional deficit imparted by the pathogenic variant (Fig. 1A). It should be noted that the EC_50_ of Org 34167 is ∼10-fold higher in the *Xenopus laevis* oocytes compared to HEK cells, explaining the high concentration used here.^13,14^ A 50:50 mix of the HCN1^M305L^ pathogenic variant RNA with HCN1^WT^ RNA was also injected into oocytes to mimic the heterozygous status of patients. ‘Heterozygous’ HCN1^WT+M305L^ channels also have reduced voltage dependence, with a proportion of channels open at more depolarised potentials, resulting in a larger instantaneous current^7,8,18^ (Fig. 1A, B). Org 34167 (100 µM) restored voltage dependence to the ‘heterozygous’ mutated channel, significantly reducing the instantaneous current (Fig. 1A, B). This is reflected in the *IV* relationship of the activation current at steady-state, where Org 34167 increased inward rectification of the steady-state activation current to resemble that exhibited by HCN1^WT^ homomeric channels (Fig. 1C). Instantaneous tail current analysis showed that the midpoint voltage (V_0.5_) was shifted 50 mV in the hyperpolarising direction to coincide closely with the typical HCN1^WT^ channel open probability (Fig. 1D). The faster activation kinetics of the ‘heterozygous’ case were only partially recovered (Fig. 1E). These data strongly support the notion that Org 34167 can restore normal channel function, overcoming the majority of the functional deficits caused by the M305L mutation, and as such could act as a precision medicine for *HCN1*-DEE.

**Figure 1.**
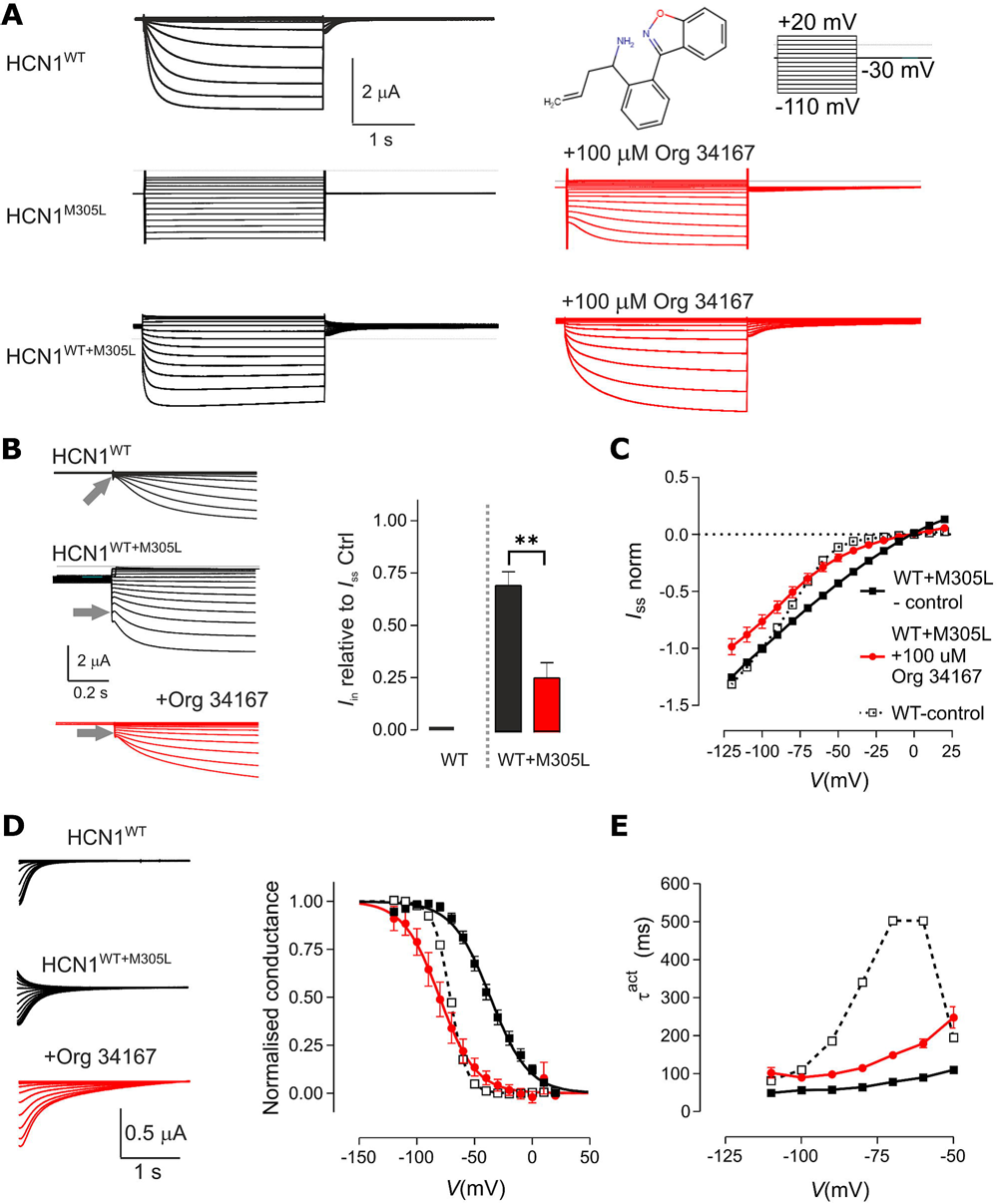
Org 34167 restores voltage dependence to the HCN1_M305L_ variant. **(A)** Representative current traces recorded from *Xenopus laevis* oocytes expressing HCN1^WT^ (top), HCN1^M305L^ alone (middle) and HCN1^WT+M305L^ co-expressed (bottom) channels in response to the activation protocol (top right), under baseline conditions (black) and in the presence of Org 34167 (100µM, red). *(insert)* Org 34167 structure. **(B)** Expanded view of current traces from (A) highlighting the instantaneous current (arrowed for the step to -100 mV). Bar graph (right) shows pooled data (n = 4) comparing the instantaneous current (I_in_) before (black) and after (red) exposure to Org 34167. **(C)** Normalised current-voltage relationship pooled (n = 4) for the steady-state activation current (I_ss_) recorded at the end of the test pulse. Filled black squares: HCN1^WT+M305L^ co-expressed oocytes at baseline; filled red circles: HCN1^WT+M305L^ co-expressed oocytes after exposure to Org 34167; unfilled squares: HCN1^WT^ expressing oocytes. **(D)** Representative tail currents for voltage steps returning to the -30 mV holding potential for HCN1^WT^ and HCN1^WT+M305L^ expressing oocytes before and after Org 34167 application. (*right*) Pooled normalised conductance vs *V*. Symbol colours as in (C). **(E)** Activation time constant (*τ*_act_) plotted against *V*. Symbol colours as in (C). **p < 0.01.

### Org 34167 reduced hyperexcitability and normalised HCN1-dependent electrophysiological properties in Hcn1^M294L^ neurons

Whole-cell patch-clamp recordings were conducted in layer V neurons from Hcn1^M294L^ mice to test the impact of Org 34167 on cellular properties (Fig. 2). I_h_-mediated sag current was measured in current clamp. Consistent with our previous results,^8^ the sag current in layer V neurons was small (Fig. 2A). Org 34167 (30 µM) significantly increased sag recorded from Hcn1^M294L^ neurons (Fig. 2A, B). As expected, Org 34167 also significantly hyperpolarised Hcn1^M294L^ layer V neurons (Fig. 2C). The AP *i-o* relationship was measured before and following the application of Org 34167, with a significant reduction in AP firing frequency observed (Fig. 2D, E, F). There was no significant shift in rheobase (Fig. 2G), which is probably a reflection of the relatively small hyperpolarising shift in membrane potential seen (Fig. 2C). There were no changes in other passive or active properties recorded (see Supplementary Table 1). Collectively, these data suggest that Org 34167 recovered I_h_ and reduced neuronal hyperexcitability caused by the Hcn1^M294L^ variant.

**Figure 2.**
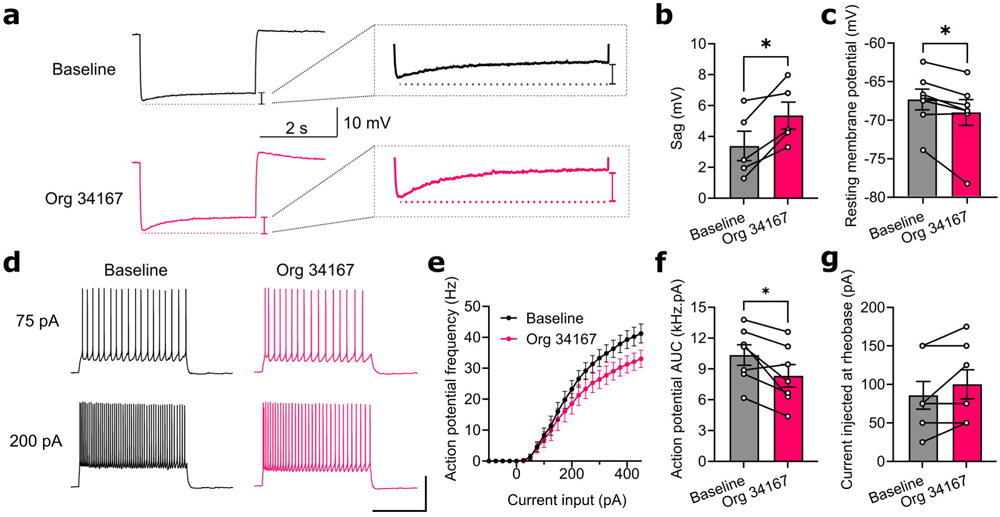
Org 34167 restores I_h_-mediated sag leading to reduced excitability of layer V pyramidal neuron excitability in the Hcn1_M294L_ mouse model of DEE. **(A)** Example sag traces obtained from a layer V pyramidal neuron from an Hcn1^M294L^ mouse at baseline (black) and following Org 34167 (30 µM, pink). **(B)** Sag is significantly increased in layer V neurons from Hcn1^M294L^ mice following administration of 30 µM Org 34167 (n = 5). **(C)** Org 34167 (30 µM) caused a significant hyperpolarisation of the resting membrane potential in Hcn1^M294L^ neurons (n = 7). **(D)** Representative AP firing patterns in response to 2 s injections of 75pA (upper) or 200pA (lower) current in layer V neurons from Hcn1^M294L^ mice at baseline (black) and following wash-on of Org 34167 (30 µM, pink). **(E)** Summary of *i-o* data showed a reduction in AP firing at larger depolarising current injections in layer V pyramidal neurons from Hcn1^M294L^ mice following Org 34167 (30 µM, n = 7). **(F)** Org 34167 (30 µM) reduced the area under the curve (AUC) of the *i-o* relationship in layer V pyramidal neurons from Hcn1^M294L^ mice (n = 7). **(G)** Org 34167 had no significant effect on rheobase of layer V pyramidal neurons from Hcn1^M294L^ mice (n = 7). *p<0.05.

### Org 34167 reduces epileptiform activity in the Hcn1^M294L^ mouse model

Changes in epileptiform ECoG spiking frequency in Hcn1^M294L^ mice provide a quantifiable measure of anti-seizure medication (ASM) efficacy with strong predictive validity of human *HCN1*-DEE patient pharmacoresponsiveness.^8,10^ Org 34167 (1 mg/kg) had no effect, while increasing the dose to 2 mg/kg significantly reduced ECoG spiking frequency (Fig. 3A, B, D). Org 34167 at both 1 mg/kg and 2 mg/kg caused changes to ECoG spectral activity, including a significant increase in power in the theta band concentrated around a peak of 6-7 Hz (Fig. 3C, E) (Supplementary Table 1). ECoG spectral activity may therefore potentially act as a biomarker of HCN channel target engagement. These data suggest that Org 34167 may be an effective ASM for *HCN1*-DEE caused by cation leak.

**Figure 3.**
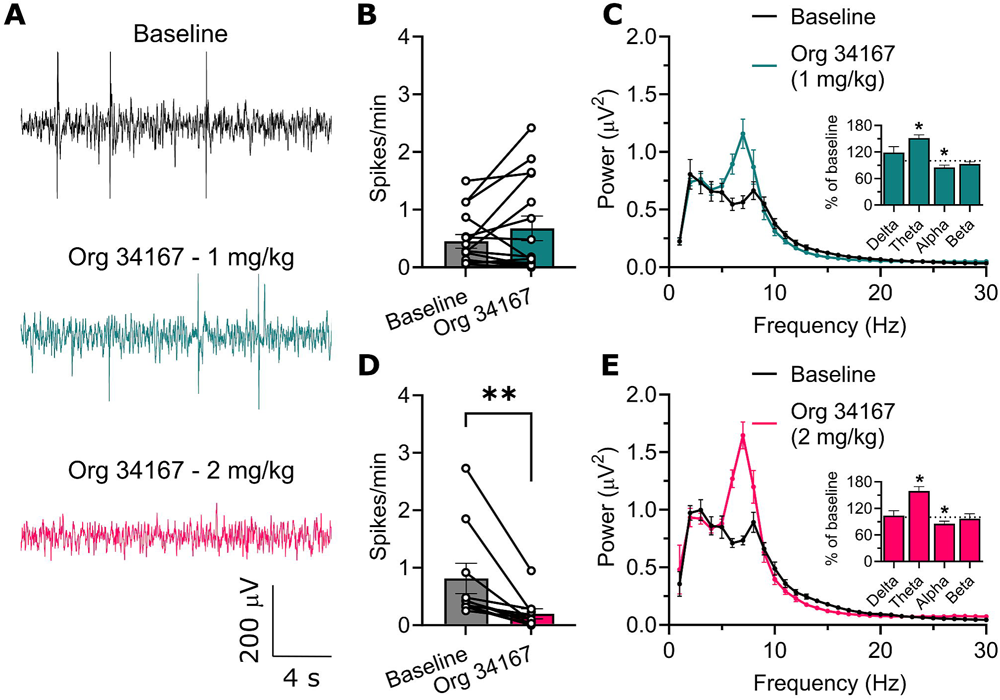
Effect of Org 34167 on ECoG spiking and spectral activity in Hcn1_M294L_ mice. **(A)** Example raw ECoG data traces from Hcn1^M294L^ mice at baseline (black), after administration of Org 34167 (1 mg/kg, blue), and after administration of Org 34167 (2 mg/kg, pink). **(B)** Org 34167 (1 mg/kg) had no significant effect on ECoG spike frequency in Hcn1^M294L^ mice (n = 15, p=0.4). **(C)** Org 34167 (1 mg/kg, n = 14) caused a significant increase in power in the theta band (p<0.0001) and a significant reduction of power in the alpha band (p=0.02), with no significant effect on delta or beta activity. **(D)** Org 34167 (2 mg/kg) caused a significant reduction in ECoG spike frequency in Hcn1^M294L^ mice (n = 10). **(E)** Org 34167 (2 mg/kg, n = 9) caused a significant increase in power in the theta band (p=0.0002) and a significant reduction of power in the alpha band (p = 0.02) with no significant effect on delta or beta activity. **p < 0.01.

### Org 34167 restores photoreceptor sensitivity in the Hcn1^M294L^ mouse retina

ERG recordings from Hcn1^M294L^ mice have revealed a significant decrease in both rod and cone photoreceptor sensitivity to light and reduced temporal processing that is recapitulated in a patient with the equivalent *HCN1* variant.^19^ This makes the ERG signature a potential biomarker of efficacy of precision therapeutic approaches in *HCN1*-DEE. As previously described, ERG traces from Hcn1^M294L^ mice in response to 2.07 log cd.s/m^2^ light pulse are attenuated and truncated when compared to Hcn1^WT^ retina (Fig. 4A, B). Interestingly, Org 34167 (2 mg/kg) had little impact on ERG responses in the Hcn1^WT^ mouse retina (Fig. 4A, B). Remarkably, for the Hcn1^M294L^ mouse, Org 34167 (2 mg/kg) restored the waveform over time so that it more closely matched Hcn1^WT^ values (Fig. 4A, B).

**Figure 4.**
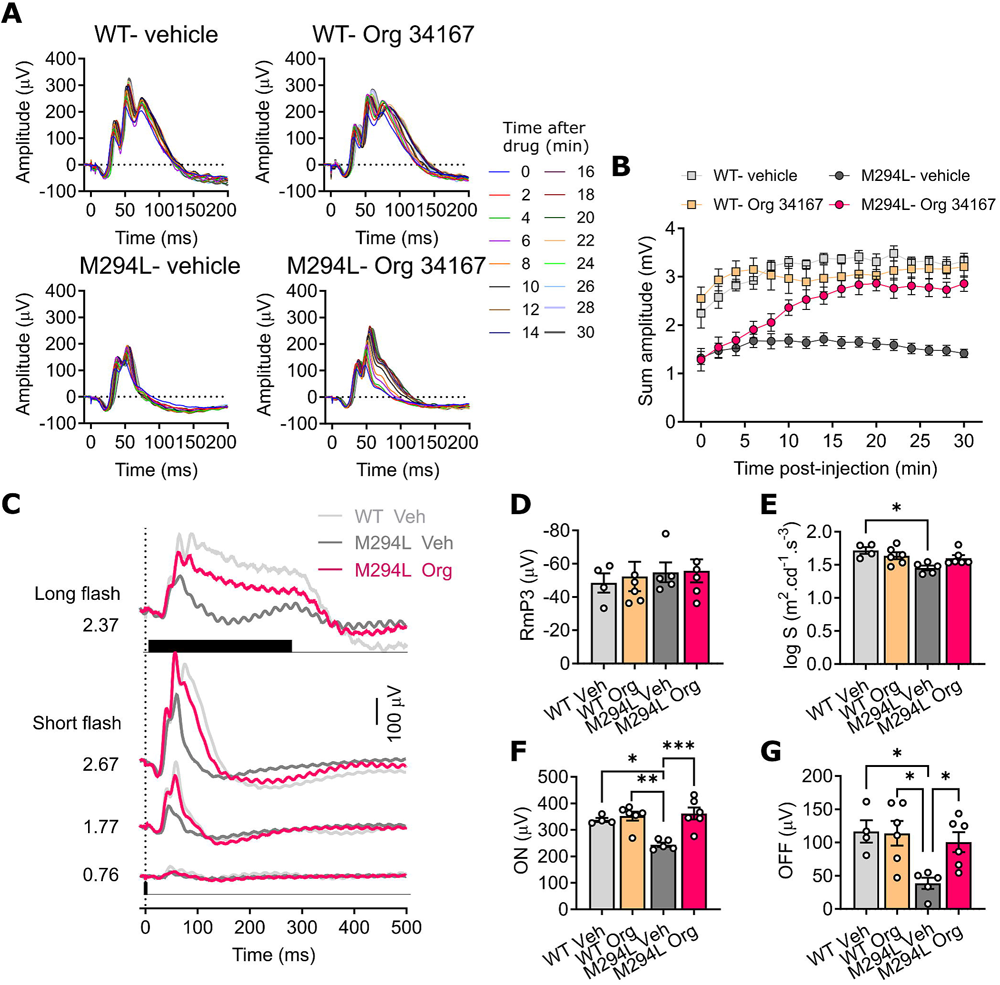
Org 34167 restores retinal function in the Hcn1_M294L_ mouse. **(A)** Group averaged ERG waveforms elicited by a 2.07 log cd.s/m^2^ light pulse recorded from Hcn1^WT^ mice (n = 4, top left), Hcn1^WT^ mice administered Org 34167 (2 mg/kg, n = 6, top right), Hcn1^M294L^ mice (n = 5, bottom left) and Hcn1^M294L^ mice administered Org 34167 (2 mg/kg, n = 6, bottom right) mice, at baseline and every 2 minutes following drug injection. **(B)** Summed amplitude under the voltage across time envelope (0-150 ms) for Hcn1^WT^ mice (grey square), Hcn1^WT^ mice administered Org 34167 (orange square), Hcn1^M294L^ mice (grey circle) and Hcn1^M294L^ mice administered Org 34167 (pink circle). **(C)** Group averaged ERG waveforms in response to short pulses at selected intensities (0.76-2.76 log cd.s/m^2^, bottom) as well as a longer 256 ms pulse (top), in Hcn1^WT^ mice administered vehicle (veh) (n = 4, light grey traces), Hcn1^M294L^ mice administered vehicle (n = 5, dark grey), and Hcn1^M294L^ mice administered Org 34167 (2 mg/kg, n = 6, pink). The black bar represents the time that light is delivered. **(D)** Photoreceptoral amplitude for the 4 groups: Hcn1^WT^ mice administered vehicle (light grey bar), Hcn1^WT^ mice administered Org 34167 (orange), Hcn1^M294L^ mice administered vehicle (dark grey) and Hcn1^M294L^ mice administered Org 34167 (pink). **(E)** Photoreceptoral sensitivity for the 4 groups. **(F)** Amplitude of post-receptoral response to light onset for the 4 groups. **(G)** Amplitude of post-receptoral response to light offset for the 4 groups. *p < 0.05; **p < 0.01; ***p < 0.001.

Furthermore, the complexity of the response to both short and long pulses was also restored (Fig. 4C). Photoreceptoral amplitudes were unchanged in Hcn1^M294L^ mice and unaffected by drug (Fig. 4D). Org 34167 almost completely recovered cone photoreceptoral sensitivity to light (Fig. 4E), as well as post-receptoral pathways that subserve responses to light increments (Fig. 4F) and decrements (Fig. 4G). Finally, temporal processing deficits were also recovered by Org 34167 (Fig. 5A, B). These data indicate reversal of deficits caused by the mutated Hcn1 channel at the retinal neuronal network scale and support the idea that Org 34167 can act as a precision therapeutic addressing not only seizures, but also comorbidities.

**Figure 5.**
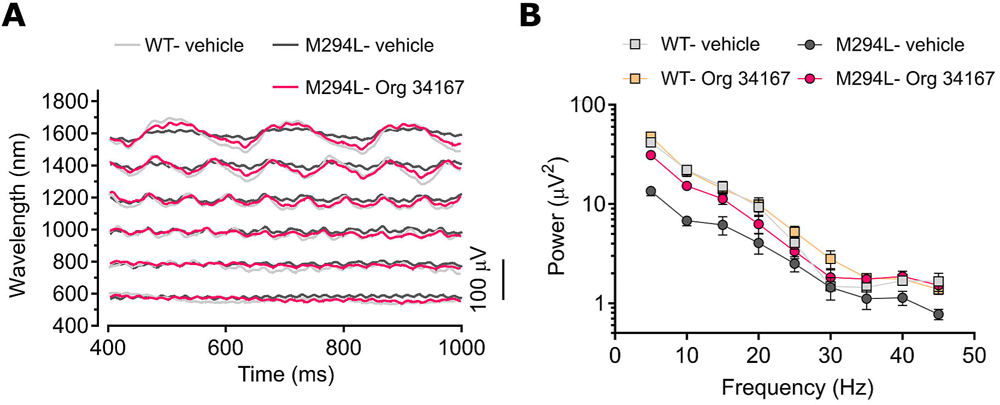
Recovery of visual response to high temporal frequencies in Hcn1_M294L_ mice by Org 34167. **(A)** Group averaged ERG waveforms from Hcn1^WT^ mice administered vehicle (n = 4, light grey), Hcn1^M294L^ mice administered vehicle (n = 5, dark grey) or Hcn1^M294L^ mice administered Org 34167 (2 mg/kg, n = 6, pink) collected at a range of stimulus frequencies (5-45 Hz), 30 minutes following drug injection. **(B)** Group average responses as a function of flicker frequency reported for Hcn1^WT^ mice administered vehicle (grey squares, n = 4), Hcn1^WT^ mice administered Org 34167 (orange squares, n = 6), Hcn1^M294L^ mice administered vehicle (grey circles, n = 4) and Hcn1^M294L^ mice administered Org 34167 (pink circles, n = 6).

### Behavioural analysis of Org 34167 in the Hcn1^M294L^ mouse model

Org 34167 has passed phase I clinical trials and is reported to be well tolerated at doses of up to 40 mg in humans.^12^ However, visual observation of Hcn1^M294L^ mice revealed that 12 out of 13 mice administered Org 34167 (2 mg/kg) displayed tremors. These ranged in severity from short-lasting and intermittent through to severe and persisting for over one hour following drug administration. Severe tremors are also observed in Hcn1^WT^ mice at this dose which is reflected in a complete lack of mobility and consistent with data previously reported^13^ (Fig. 6A). Interestingly, Org 34167 (2 mg/kg) did not reduce locomotion with the Hcn1^M294L^ mice maintaining their hyperactive state (Fig. 6A). Org 34167 (2 mg/kg) did reduce the time Hcn1^M294L^ mice spent on the rotarod, although this was not significantly different from saline injected Hcn1^WT^ mice (Fig. 6B). Org 34167 (2 mg/kg) normalised performance on the elevated plus maze back to Hcn1^WT^ levels, although these data are hard to interpret in the context of potentially improved vision and drug toxicity (Fig. 6C). Collectively these data suggest that an efficacious Org 34167 dose causes behavioural side-effects, although the Hcn1^M294L^ mice tolerate the drug better than Hcn1^WT^ mice.

**Figure 6.**
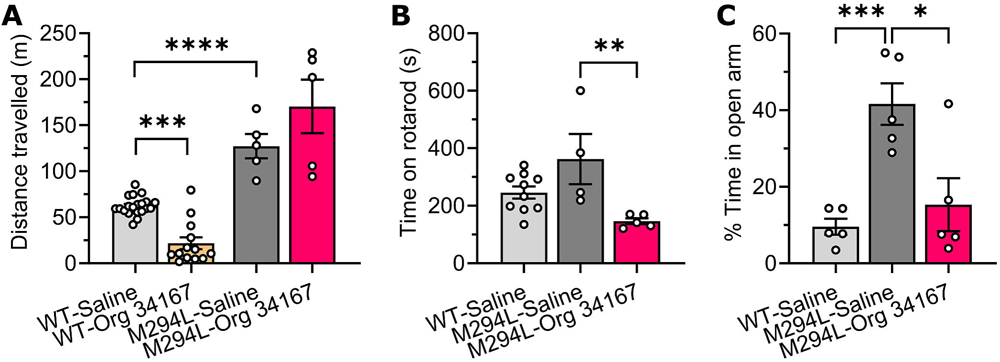
Behavioural analysis of Org 34167 in the Hcn1_M294L_ mouse model. **(A)** Hcn1^M294L^ mice travelled significantly further than Hcn1^WT^ littermates in the open field assay (n = 5 Hcn1^M294L^, n = 20 Hcn1^WT^). Org 34167 (2 mg/kg) caused a significant reduction in the distance travelled by Hcn1^WT^ mice (n = 13), whereas it had no significant effect on the distance travelled by Hcn1^M294L^ mice (n = 5). **(B)** Hcn1^M294L^ mice administered Org 34167 (2 mg/kg, n = 5) fell from the rotarod sooner than Hcn1^M294L^ mice administered vehicle control (n = 4). Org 34167 (2mg/kg) caused Hcn1^M294L^ mice to fall sooner from the rotarod than Hcn1^WT^ mice (n = 10). **(C)** Hcn1^M294L^ mice spent significantly longer in the open arms of the elevated plus maze than Hcn1^WT^ littermates. Org 34167 (2 mg/kg) normalised performance of Hcn1^M294L^ mice on the elevated plus maze. (n = 5 for all groups). *p < 0.05; **p < 0.01; ***p < 0.001.

### Org 34167 reduces cation leak caused by other *HCN1*-DEE variants

We tested the efficacy of Org 34167 against 5 additional *HCN1*-DEE variants expressed in *Xenopus* oocytes (Fig. 7). As previously described, each of the tested variants exhibited cation leak, a common biophysical feature of *HCN1*-DEE.^7^ Org 34167 significantly reduced cation leak in all 5 additional *HCN1*-DEE variants (Fig. 7A, B). This consistent effect underscores the potential utility of Org 34167 as a therapeutic strategy in *HCN1*-DEE more broadly.

**Figure 7.**
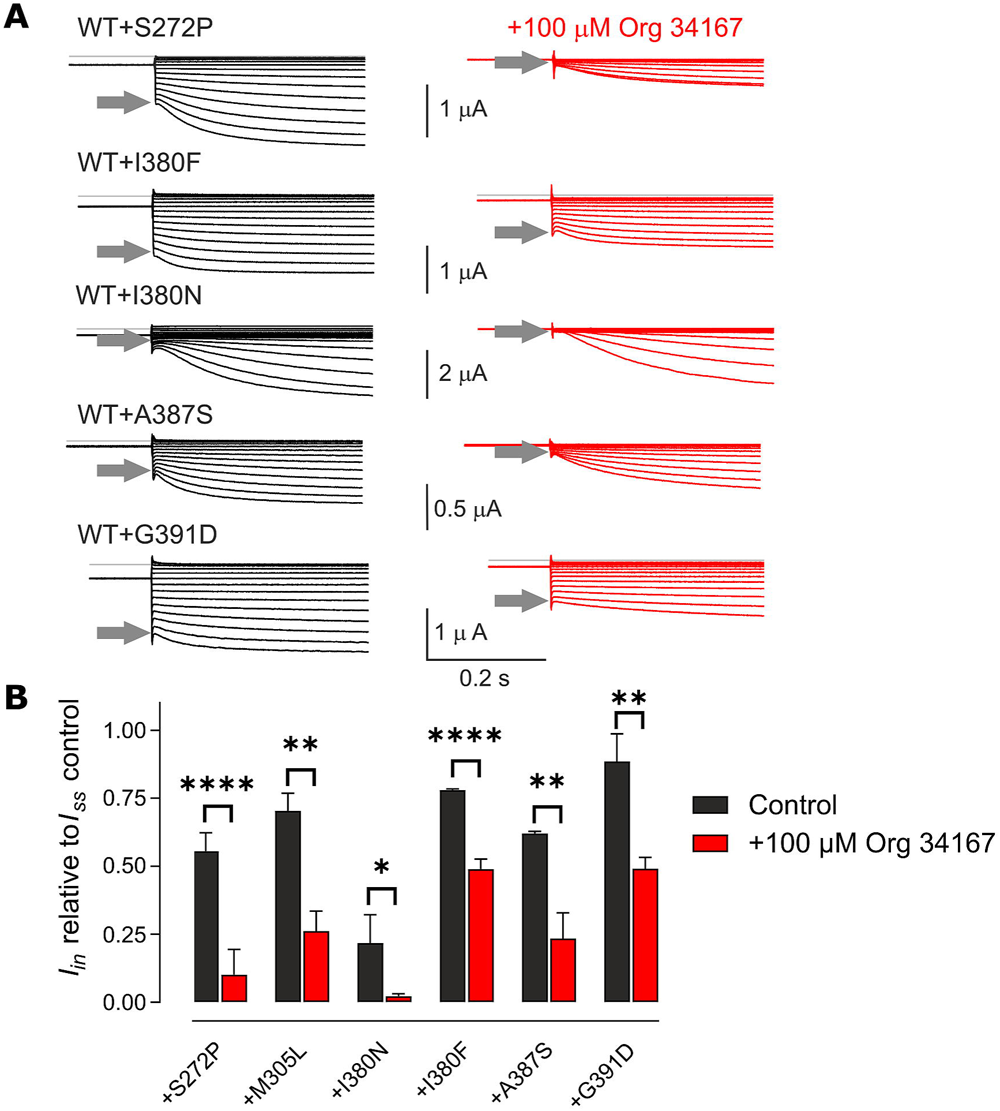
Org 34167 significantly reduces the instantaneous (leak) current in DEE variant channels. **(A)** Representative current traces recorded from oocytes expressing ‘heteromeric’ DEE variant channels (50:50 mixture of cRNA for HCN1^WT^ and the respective variant) in response to the voltage step protocol shown in Fig. 1A. The effect of Org 34167 exposure on the instantaneous current (arrowed for step -100 mV) is clearly seen. **(B)** Pooled data (n ≥ 3) for the DEE variant channels showing *I*_in_ at -100 mV normalised to the steady-state current at -100 mV at baseline (black) and after exposure to 100 μM Org 34167 (red). *p < 0.05; **p < 0.01; ****p < 0.0001.

## Discussion

*HCN1*-DEE is a devastating disease that is characterised by severe seizures, intellectual disability, and other comorbidities.^4^ Treatment is limited to ASMs, with emerging evidence that sodium valproate is the most effective in *HCN1*-DEE.^2,3,5,8,10,20^ However, *HCN1*-DEE patients remain mostly refractory to standard ASMs.^2,7^ Comorbid states need to be treated independently, with parent reports suggesting that these are also poorly controlled with standard of care therapeutics. This highlights an urgent need to develop precision therapeutics for *HCN1*-DEE. Here we test the brain penetrant HCN channel modulator, Org 34167, in the Hcn1^M294L^ mouse model of *HCN1*-DEE. Org 34167 reduced neuronal epileptiform activity and recovered retinal light sensitivity and temporal responsivity, suggesting it could potentially treat both seizures and comorbidities. Org 34167 also reduced cation leak caused by other *HCN1*-DEE pathogenic variants suggesting that it may be more broadly used in this disease. At effective doses, Org 34167 caused tremors in Hcn1^M294L^ mice, although these seem less severe than in Hcn1^WT^ mice.

Org 34167 successfully completed phase I trials for depression and was reported to be well tolerated at doses as high as 40 mg.^12^ The side effects for humans included dizziness, impaired concentration, nausea and vomiting, sleeping disturbances, vertigo, fatigue, headache, palpitations and visual disturbances.^12^ However, our data suggests that tremors emerge at therapeutic doses in the Hcn1^M294L^ mouse. Tremors are likely to be mediated by modulation of HCN2 channels, given that *Hcn2* knockout mice are reported to have severe tremors,^21^ whereas *Hcn1* and brain-specific *Hcn4* knockout mice are tremor-free.^22–24^ Interestingly, tremors are not mentioned as a side-effect in humans,^12^ suggesting there may be a disconnect between species with regard to the side-effects experienced.

HCN4 channel block is thought to underlie the bradycardic effect of the clinically used, peripherally restricted, broad-spectrum HCN channel inhibitor, ivabradine.^25^ As expected, the broad-spectrum HCN channel modulation of Org 34167 does cause heart rate reduction in mice, which may translate to be a confound for clinical use.^13^ However, ivabradine only causes a self-limiting bradycardia and is otherwise extremely well tolerated.^25–27^ Furthermore, Org 34167 concentrates in the brain (blood plasma: brain ratio ∼17) which will limit peripheral side effects. ^13^ Nevertheless, new compounds with less efficacy against HCN2 and HCN4 channels would be predicted to have a larger therapeutic window, potentially reducing the incidence of tremor and further reducing the impact on the heart.

An increase in the proportion of the HCN1 channel-mediated instantaneous current relative to the total current is a useful biomarker for a significant proportion of *HCN1*-DEEs.^7^ Remarkably, Org 34167 recovered much of the voltage dependence of the ‘homomeric’ and ‘heteromeric’ HCN1^M305L^ mutated channels, restoring them back to HCN1^WT^ levels. Similar recoveries can be seen in channels comprising other HCN1 cation leak variants, which suggests that patients carrying these may also benefit from Org 34167 treatment. Equally, *HCN1* variants that cause depolarising shifts in the voltage of activation could potentially benefit from the hyperpolarising effects of Org 34167. However, Org 34167 does impact distinct *HCN1* variants to differing extents, as reflected in the variability in the relative reduction in instantaneous current (Fig. 7B). As such, although our prediction is that Org 34167 will be effective in these cases, the extent of any benefits observed may be *HCN1* variant specific, with additional preclinical studies needed to test this idea.

Our data has broad implications for the treatment of DEEs, especially those caused by pathogenic variation in genes encoding ion channels. Screening approved drugs against specific biophysical markers in mutated channels may reveal repurposing candidates. Importantly, as highlighted here, the impact on WT channels may not necessarily be a predictor of efficacy. The fact that Org 34167 restores voltage sensitivity to the mutated HCN1 channel is distinct to the inhibitory action seen in the WT situation. Furthermore, small molecules are likely to have a better biodistribution than many of the genetic therapeutics currently under development. This means that they will potentially address comorbidities that are associated with different organ systems where mutated channels are expressed. The potential impact of this is highlighted by the restoration of retinal function in the Hcn1^M294L^ mouse.

In summary, we provide strong preclinical evidence that Org 34167 or similar small molecule HCN modulators could act as precision medicines for *HCN1*-DEE. The repurposing of Org 34167 for *HCN1*-DEE should be considered, although evidently a full toxicological profile of Org 34167 and information on pharmacokinetics and formulation would be needed. Small molecules developed to have HCN1 channel selectivity over HCN2 and HCN4 channels are likely to have wider therapeutic windows and should be pursued for future use in *HCN1*-DEE.

## Funding and Acknowledgements

The Florey Institute of Neuroscience and Mental Health acknowledges the strong support from the Victorian Government and in particular the funding from the Operational Infrastructure Support Grant. This work was also made possible through the Australian Government National Health and Medical Research Council (NHMRC) independent research Institute Infrastructure Support Scheme (IRIISS). We acknowledge the support of an NHMRC Program Grant (10915693) to CAR. LEB acknowledges the support of an Australian Government Research Training Program Scholarship. The Hcn1^M294L^ mouse line was produced by the Monash University node of the Australian Phenomics Network (APN). The APN is supported by the Australian Government through the National Collaborative Research Infrastructure Strategy (NCRIS) Program.

## Author contributions

CAR, BVB and ICF designed research studies. LEB, CEM, DZ, JS, ICF and BVB all completed experiments that contributed to the data presented in the manuscript. LEB, CEM, DZ, JS, MSS, ICF and BVB all completed analysis of data presented in the manuscript. CAR and LEB wrote the first draft of the manuscript. All authors contributed to the revision of the manuscript. CAR and BVB funded the project.

## Supplementary material

Supplementary material is available at Brain online.

### Abbreviations

aCSF: Artificial cerebrospinal fluid
AP: Action potential
ASM: Anti-seizure medication
AUC: Area under the curve
DEE: Developmental and epileptic encephalopathy
ECoG: Electrocorticography
ERG: Electroretinography
HCN: Hyperpolarisation-activated, Cyclic Nucleotide-gated channel
HEK: Human embryonic kidney
I_h_: Hyperpolarisation-activated current
i.p.: Intraperitoneal
IV: Current-voltage
SEM: Standard error of the mean
V_0.5_: Half-maximal activation voltage
WT: Wild-type

