## Supplementary Material for "A precision medicine approach for *HCN1* Developmental and Epileptic Encephalopathy"

**Supplementary Table 1.** Summary of numerical data from electrophysiology experiments presented in Figure 2, electrocorticography experiments presented in Figure 3, and behavioural experiments presented in Figure 4. All data are presented as mean ± SEM. Data for Figure 2 were analysed using a paired two-tailed Student's *t* test. Data for Figure 3 were analysed using a Wilcoxon test. Data for Figure 4 were analysed using an unpaired two-tailed Student's *t* test.

**Figure 2**

| **Parameter** | **Baseline** | **Post ORG 34167** | **P-value** |
|---|---|---|---|
| Sag (mV) (Fig. 2B) | 3.4 ± 1.0<br>N = 5 | 5.4 ± 1.9<br>N = 5 | 0.02 |
| RMP (mV) (Fig. 2D) | -67.4 ± 1.3<br>N = 7 | -69.1 ± 1.7<br>N = 7 | 0.02 |
| Rheobase (pA) (Fig. 2G) | 85.7 ± 18<br>N = 7 | 100 ± 19<br>N = 7 | 0.1 |

**Figure 3**

| **Parameter** | **Baseline** | **Post ORG 34167** | **P-value** |
|---|---|---|---|
| Spike frequency at 1mg/kg ORG 334167 (Fig. 3B) | 0.5 ± 0.1<br>N = 15 | 0.7 ± 0.2<br>N = 15 | 0.37 |
| Spike frequency at 2mg/kg ORG 334167 (Fig. 3D) | 0.8 ± 0.3<br>N = 10 | 0.2 ± 0.1<br>N = 10 | 0.002 |

**Figure 4**

| **Parameter** | **WT + saline** | **WT + ORG 34167** | **Hcn1$^{M294L}$ + saline** | **Hcn1$^{M294L}$ + ORG 34167** | **P-value** |
|---|---|---|---|---|---|
| Locomotor distance travelled (m) (Fig. 4A) | 62 ± 8<br>N = 5 | 5 ± 1<br>N = 5 | | | 0.0001 |
| | | | 127 ± 13<br>N = 5 | 170 ± 29<br>N = 5 | 0.2 |
| | 62 ± 8<br>N = 5 | | 127 ± 13<br>N = 5 | | 0.003 |
| Elevated Plus Maze Percentage time in open arm (%) (Fig. 4B) | | | 42 ± 5<br>N = 5 | 15 ± 7<br>N = 5 | 0.02 |
| | 10 ± 2<br>N = 5 | | 42 ± 5<br>N = 5 | | 0.0006 |
| | 10 ± 2<br>N = 5 | | | 15 ± 7<br>N = 5 | 0.4 |
| Rotarod time to fall (s) (Fig. 4C) | | | 362 ± 87<br>N = 4 | 146 ± 10<br>N = 5 | 0.03 |
| | 248 ± 36<br>N = 5 | | 362 ± 87<br>N = 4 | | 0.2 |
| | 248 ± 36<br>N = 5 | | | 146 ± 10<br>N = 5 | 0.03 |

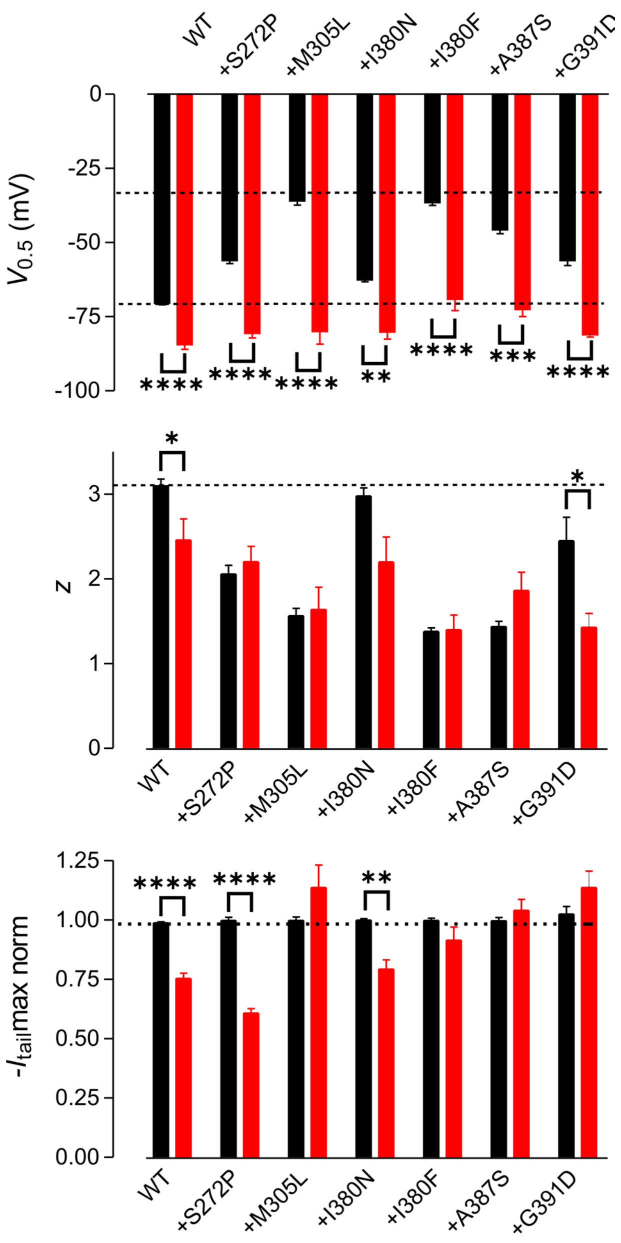


**Supplementary Figure 1. Impact of Org 34167 on the biophysical parameters of DEE variant channels.**

Pooled data ($n \geq 3$) for the DEE variant channels comparing the baseline (black) and after exposure to 100 µM Org 34167 (red) on voltage for half maximal activation (top), slope factor (z, middle), and normalised maximal tail current (bottom). *$p < 0.05$; **$p < 0.01$; ****$p < 0.0001$.
